# Improving parameter inference by resolving Bayesian prior ambiguity via multi-dataset analysis: Application to isothermal titration calorimetry

**DOI:** 10.1101/2025.07.09.663792

**Authors:** Lisa Otten, Douglas R. Walker, Elisar J. Barbar, Daniel M. Zuckerman

## Abstract

Isothermal titration calorimetry (ITC) is a powerful technique for probing biomolecular interactions. However, accurate determination of binding parameters—such as enthalpy and free energy—as well as associated uncertainties can be hindered by noise and concentration variability. Notably, the mathematical ambiguity surrounding analyte concentrations in standard binding models intrinsically limits the precision with which binding parameters, particularly binding enthalpies, can be determined. Here, we present a Bayesian pipeline that resolves this ambiguity by combining two key strategies: simultaneous analysis of multiple ITC datasets and a hierarchical Bayesian treatment of analyte concentration priors. This dual approach not only lifts the degeneracy inherent in single-dataset studies but also removes an ambiguity typically present in Bayesian analysis by self-consistently refining concentration estimates, ensuring optimal joint inference of binding parameters and concentrations. Using modern Monte Carlo techniques enables our pipeline to provide robust posterior sampling for more than 10 datasets and 40 total parameters. We validate the approach with synthetic ITC datasets for single- and multi-site binding models and apply it to experimental data, including 14 datasets for 1:1 binding of Mg(II) to the chelator EDTA and multiple datasets of the hub protein LC8 with diverse binding partners. This work serves as a foundation for improving the precision of binding constants using multiple ITC datasets, while providing a systematic framework for assessing the reliability of experimental concentration estimates, paving the way for more accurate biomolecular interaction studies.

## 1 Introduction

Biophysical experiments generate data that must be interpreted through the lens of quantitative models. These models rely on parameters that connect experimental observables to underlying physical processes. This study applies and compares different Bayesian approaches for parameter estimation from multiple biophysical data sets, i.e., replicate experiments, as exemplified by the case of isothermal titration calorimetry (ITC). Here, ITC exemplifies the broader challenges of parameter inference, namely, the complex interplay between nuisance parameters, such as analyte concentrations, and critical parameters of interest, including binding affinities and enthalpies. These challenges emphasize the need for advanced analytical techniques capable of addressing correlation structure and delivering robust error estimates. While this study centers on ITC to provide practical guidance for practitioners addressing these challenges in biophysical and chemical measurements, the methodologies and insights presented are likely applicable to analogous problems across other fields.

ITC is a powerful and widely used technique for investigating molecular interactions in solution, with applications across numerous scientific disciplines [1]. ITC directly measures the heat change Δ*Q* in a solution resulting from compositional changes during titration [2, 3]. This enables the determination of key thermodynamic parameters, such as binding affinities, enthalpies, and entropies.

Consequently, ITC is employed to investigate a wide range of binding interactions, including protein-protein [4], protein-ligand [5], and protein-RNA systems [6, 7]. ITC can probe 1:1 binding and also more complex processes such as competitive [8, 9] and cooperative binding [10, 11]. Despite its versatility and utility, ITC data analysis presents multiple challenges that impact the accuracy in determining binding parameters and their error estimates under real-world experimental conditions.

In recent years, Bayesian inference (BI) [12] has found growing utility across diverse areas of biophysics as a powerful framework for interpreting noisy and complex experimental data. Notable applications include ion channel modeling [13, 14, 15], single-molecule trajectory analysis [16, 17, 18, 19], and the modeling of biochemical networks [20, 21]. Likewise, BI has emerged as a powerful tool to improve ITC analysis [22, 23, 24, 25, 26, 11]. In essence, BI is a statistical framework that generates a “posterior” probability distribution of model parameters based on the dataset(s) and prior information, such as constraints, on the model parameters.

While computationally more expensive than alternative frequentist methods, BI offers several advantages. It provides a comprehensive posterior distribution that contains most likely parameter estimates, uncertainties, and the full correlation structure between all binding and nuisance parameters. Combined with the explicit modeling of analyte concentration errors [22] and the ability to implement modular frameworks for analyzing multiple datasets [27] and distinguishing between different binding models [25], BI enables more accurate parameter estimation while accommodating complex experimental designs and prior knowledge. However, BI’s reliance on user-defined prior distributions is sometimes viewed as a limitation, as the choice of priors can influence the results and requires careful consideration to avoid bias.

In the case of ITC, the experimental workflow converts differential power measurements into integrated heat values, which form the foundation for subsequent analysis [2]. Thermodynamic parameters are then determined by fitting binding heat models based on the mass action law to the integrated heat. Unfortunately, the intrinsic information content of individual isotherms is limited by an inherent mathematical degeneracy in binding models that results in a natural ambiguity of thermodynamic parameters and analyte concentrations [11] making the accurate determination of parameters more challenging.

Standard parameter estimation protocols, such as those implemented in the Origin software package [28], rely on nonlinear least squares fitting to estimate the association constant *K*_*a*_, the binding enthalpy Δ*H*, and the stoichiometry *n*. However, these methods have limitations, particularly in assessing parameter correlations and uncertainties. Notably, while errors in the titrand concentration are partially addressed by allowing the stoichiometry *n* to vary as a non-integer parameter, the titrant concentration is generally assumed to be precisely known. In practice, titrant concentration errors of 10–20% are common and challenging to estimate [29], leading to downstream effects on the precision and reliability of the inferred parameters [30]. While not typically performed, explicit treatment of titrant concentration errors in nonlinear least squares fitting is possible. One example of this is the SEDPHAT package [31], which not only incorporates a more sophisticated error model, but also allows for the global analysis of multiple datasets improving parameter inference by leveraging information across experiments.

In this manuscript, we build on previous advancements in BI for ITC [22], multi-dataset analysis [23], and the mathematical degeneracy in ITC binding models [11] to propose a robust data analysis protocol for multiple isotherms. This protocol combines multi-dataset BI with a hierarchical approach that leverages the set of binding parameters common to all datasets to dynamically update and refine analyte concentration priors. This dual strategy addresses the issue of mathematical degeneracy while self-consistently accounting for concentration measurement errors, ultimately enabling more accurate determination of thermodynamic parameters. Using modern Monte Carlo techniques [32] enables our pipeline to provide robust posterior sampling for more than 10 datasets and 40 total parameters. While tailored to ITC, many aspects of the pipeline are readily transferable to other biophysical problems.

Our modular analysis pipeline, implemented as a flexible Python package [33], supports a broad range of binding models, including simple monomeric binding as well as more complex scenarios such as cooperative and competitive binding. To validate the approach, we analyze both synthetic and experimental ITC datasets, including 14 datasets on the 1:1 binding of Mg(II) to the chelator EDTA, previously studied individually in Ref. [22]. Additionally, we compare the hierarchical pipeline against other Bayesian methods, including an empirical evaluation and a standard approach that uses fixed analyte concentration priors. Finally, we outline how synthetic data can complement experimentation to design more informative experiments.

## 2 Methods

In this study, we employ a Bayesian framework to analyze both synthetic and experimental data to evaluate parameter precision, optimize the modeling of experimental uncertainties, and explore different experimental conditions. Synthetic datasets allow for cost-effective, systematic testing across a broad range of parameter regimes, with the advantage of a known ground truth. This facilitates method development and the design of more informative experiments. To address limitations inherent in synthetic systems—such as oversimplification or omission of real-world noise—we complement this analysis with experimental data. Specifically, we examine the well-characterized 1:1 complexation reaction between Mg(II) and EDTA [22], as well as binding of the dimeric hub protein LC8 with multiple binding partners, which presents additional complexity due to cooperative and multivalent interactions.

### 2.1 Bayesian Inference

BI is a statistical approach used to determine the posterior distribution of model parameters by updating prior assumptions with observed data [12]. Prior distributions represent initial beliefs about parameter values before any data is considered. As more data is incorporated, the influence of these priors diminishes. The resulting posterior distribution provides valuable insights, including parameter estimates, uncertainty quantification through credibility intervals, and correlations that reveal how parameters vary together.

In this framework, a model is used to predict observables based on a set of parameters ***θ***, and Bayes’ theorem is applied to update the probability of those parameters in light of observed data ***D***

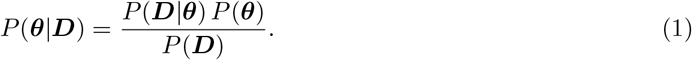

The individual terms correspond to the following:

- *P* (***θ*** |***D***) is the posterior probability, representing the updated belief about the model parameters ***θ*** after observing the data ***D***.
- *P* (***D***| ***θ***) is the likelihood, describing the probability of the observed data ***D*** given the model parameters ***θ***.
- *P* (***θ***) is the prior probability, encoding prior knowledge or assumptions about the model parameters ***θ*** before observing the data.
- *P* (***D***) is the marginal likelihood, acting as a normalizing constant to ensure the posterior is a valid probability distribution.

The dataset ***D*** ≡ {*y*_1_, *y*_2_, …, *y*_*N*_} contains the observed measurements, and the model parameters ***θ*** determine the predictions ***y***_pred_. The parameter set may also include nuisance parameters such as noise variances or instrumental bias necessary for computing the likelihood. These are detailed below.

#### 2.1.1 Likelihood

We adopt a standard likelihood function assuming Gaussian observational noise

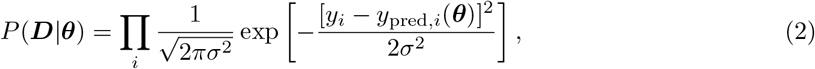

where *σ* is the standard deviation of the measurement noise. This form assumes that each observation is independent and identically distributed such that the likelihood function factorizes for each data point. Likewise, the combined likelihood for multiple datasets is given by the product of the individual likelihood functions [27].

#### 2.1.2 Priors

In its simplest form, the prior probability *P* (***θ***) factorizes across parameters

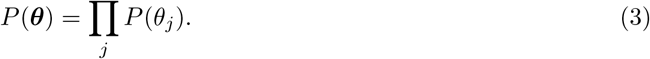

In the absence of prior information, broad uniform priors are used. The prior for the noise standard deviation *σ* is also chosen uniformly over a practical range. When available, prior information from previous experiments, independent measurement values, or theoretical constraints can be incorporated to improve inference quality.

#### 2.1.3 Obtaining the Posterior

Because the posterior distribution for a complex system is very high dimensional, we study it using a set of samples (***θ***_1_, ***θ***_2_, …) distributed according to (1). In practice, we generate samples using a recently developed preconditioned Monte Carlo approach implemented in the Python package pocoMC [32] to sample from the posterior distribution. This method addresses the limitations of traditional Markov chain Monte Carlo approaches [34, 35], which may prove challenging to converge in systems characterized by high-dimensional parameter spaces and complex sampling landscapes, akin to rough energy surfaces. The pocoMC framework combines an annealing approach with normalizing flows, enhancing convergence efficiency and enabling the convenient estimation of marginal likelihoods, which facilitates straightforward model comparison.

#### 2.1.4 Reweighting of Posterior

As part of our analysis, we examine changes in the posterior distribution resulting from different choices of priors, and these can be computed from a MC run based on a single prior via reweighting. This technique is a form of importance sampling, where the original posterior *P* (***θ***| ***D***) ∝ *P* (***D*** |***θ***) *P* (***θ***) acts as the proposal distribution, and the new posterior *P* (***θ*** |***D***)^*′*^ ∝ *P* (***D***| ***θ***) *P* (***θ***)^*′*^ is the target. The importance weight of each sample is given by

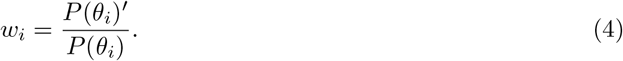

After normalizing these weights, they can be used to compute expectations under the new posterior. Reweighting is particularly useful for sensitivity analysis and model comparison when prior assumptions are varied. However, it is most reliable when the original and new prior are similar—otherwise, the resulting high variance in weights can degrade the effective sample size and as a result the approximation quality.

### 2.2 Binding Model

To infer thermodynamic binding parameters from ITC datasets, as illustrated in Fig. 1**(a)**, we generate isotherm curves using binding heat models based on the law of mass action. These predicted curves are then evaluated using Eq. (2) to calculate the likelihood of a given set of thermodynamic parameters. For simple 1:1 binding, the integrated heat is a function of [22]

**Figure 1:**
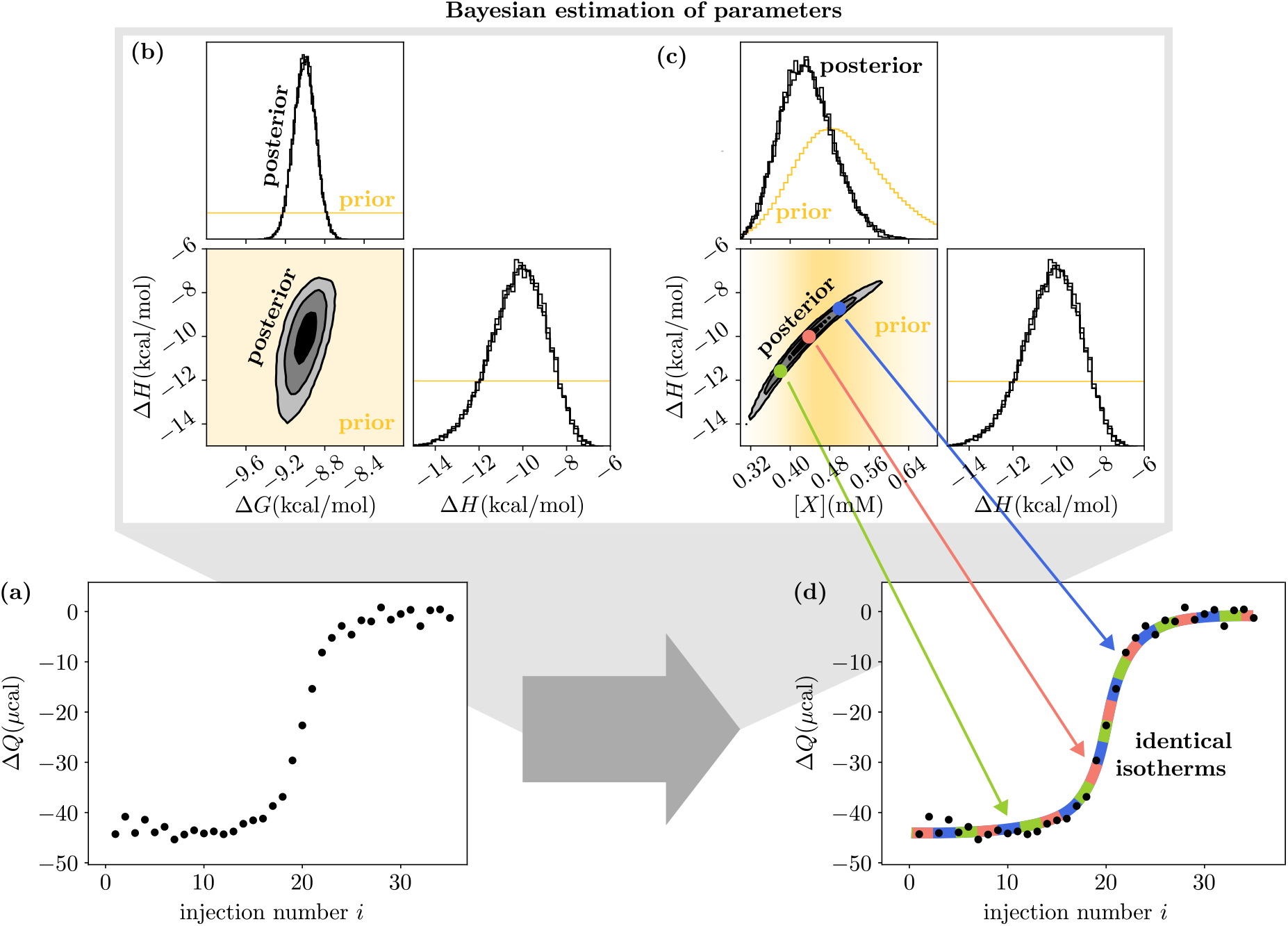
Bayesian inference for ITC and the degeneracy problem. **(a)** Example synthetic isotherm for a simple 1:1 binding model showing the integrated heats Δ*Q* per injection *i*. **(b**,**c)** Corner plots displaying the Bayesian prior and three replicates of the Bayesian posterior for the isotherm shown in (a). The posterior results show a strong correlation between enthalpy Δ*H* and titrant concentration [*X*], due to a mathematical degeneracy in the binding heat model that yields exactly equivalent isotherms for an infinite set of parameters characterized by Eq. (13). The free energy Δ*G* is less susceptible to this degeneracy. **(d)** Illustration of multiple sets of parameters leading to identical isotherm fits.

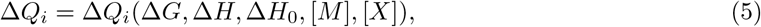

where Δ*Q*_*i*_ represents the model-predicted integrated heat at injection *i* = 1, …, *N* for an ITC curve in the absence of instrumental noise. This function depends on the following parameters:

- Δ*G*, the free energy of the microscopic binding steps with the corresponding dissociation constant *K*_*d*_ = exp(Δ*G/RT*);
- Δ*H*, the enthalpy of binding;
- Δ*H*_0_, an enthalpic correction term to account for heat of dilution, stirring per injection, buffer mismatch, and other effects that may apply a constant shift to binding heat;
- [*M*] and [*X*], the initial concentrations of the titrand in the cell and of the titrant in the syringe, respectively.

Following the identical-sites binding model implemented in Origin [28], and assuming a 1:1 stoichiometry, the integrated heat per injection is given by [36]

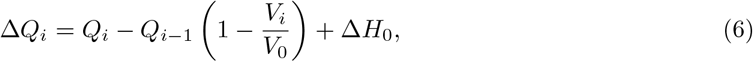

where *V*_*i*_ is the injection volume and *V*_0_ is the cell volume. The total heat *Q*_*i*_ follows the standard quadratic binding equation derived from the mass action law [2]

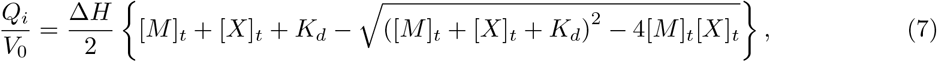

where [*M*]_*t*_ and [*X*]_*t*_ refer to the total concentration of titrand and titrant in the cell after titration step *i*.

While the quadratic expression in Eq. (7) is specific to the simple 1:1 binding model, Eqs. (5) and (6) can be readily extended to describe more complex binding with *N*_*S*_ *>* 1 distinct binding stages as well as *N*_*M*_ titrand(s) and *N*_*X*_ titrant(s). In this generalized framework, the free energy Δ*G* and enthalpy Δ*H* become vectors **Δ*G*** and **Δ*H*** of length *N*_*S*_, with each element representing the standard free energy or enthalpy associated with the formation of a particular species *S*_*j*_. Analogously, the scalar concentrations [*M*] and [*X*] are replaced by vectors [***M***] and [***X***] of length *N*_*M*_ and *N*_*X*_. Accordingly, the integrated heat released or absorbed during titration step *i* generalizes to

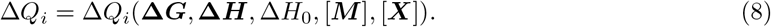

The total heat *Q*_*i*_, as used in Eq. (6), is computed from the equilibrium concentrations of the binding species, which in turn are obtained by numerically solving the following system of nonlinear equations derived from the law of mass action

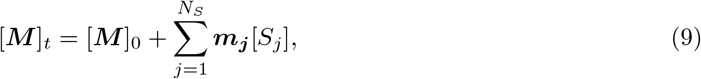

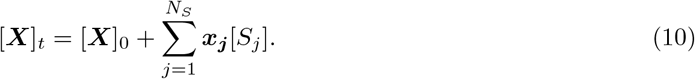

Here, [***M***]_0_ and [***X***]_0_ denote the concentrations of free titrand(s) and titrant(s) after titration step *i* to be determined. The equilibrium concentration of each complex species *S*_*j*_ is given by

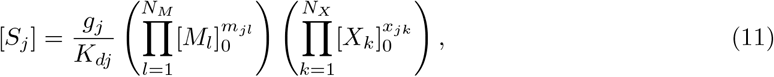

where *g*_*j*_ is a degeneracy factor accounting for indistinguishable microscopic configurations of binding. The stoichiometric vectors ***m***_***j***_ and ***x***_***j***_ specify the number of titrand and titrant molecules, respectively, involved in the formation of species *S*_*j*_. Finally, the heat is calculated as the enthalpy-weighted sum of species concentrations

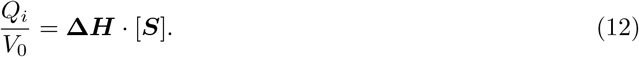

#### 2.2.1 Mathematical Degeneracy

As detailed in Ref. [11], accurate analysis of ITC data is fundamentally limited by a mathematical degeneracy in the relationship between thermodynamic parameters and analyte concentrations. As a result, ITC data describing a 1:1 binding model inherently allows for the estimation of only the ratio of titrant to titrand concentrations, while the absolute values of these concentrations remain ambiguous. This degeneracy propagates into the thermodynamic parameters, leading to strong correlations and broad distributions, where multiple parameter sets can equally describe experimental data. This invariance holds not only for the 1:1 binding model but also translates to more complex multi-site models, emphasizing the need for careful consideration of parameter interdependence when modeling ITC data.

In general, the degeneracy can be demonstrated by introducing a scaling factor *α* to apply the transformation

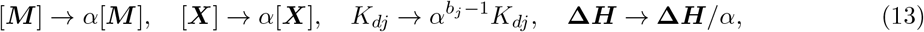

with *b*_*j*_ the number of binding partners required to form the complex species *S*_*j*_ that relates to the stoichiometric vectors of Eq. (11) via

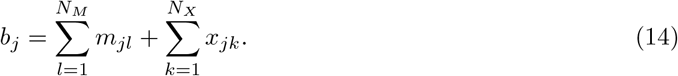

For the 1:1 binding model with *b*_*j*_ = 2, it is straightforward to verify that the transformations in Eq. (13) leave Eq. (7) unchanged. More generally, these transformations imply that the solutions for the concentrations of free titrand(s) and titrant(s), [***M***]_0_ and [***X***]_0_, are simply scaled by *α* such that [***S***] → *α*[***S***]. This scaling behavior can be confirmed by direct substitution into the transformed concentration-sum Eqs. (9) and (10). Inserting the transformed value into Eq. (12), cancels *α*, demonstrating the degeneracy in a more general setting.

As illustrated in Fig. 2, analyzing multiple ITC datasets of the same binding partners can resolve this degeneracy and narrow down both thermodynamic parameters as well as concentration estimates. This is possible because multiple datasets and their respective concentrations are coupled via the joint set of thermodynamic parameters. As the thermodynamic parameters are narrowed down by the increased amount of datasets, this narrowing is directly translated to the concentration estimates.

**Figure 2:**
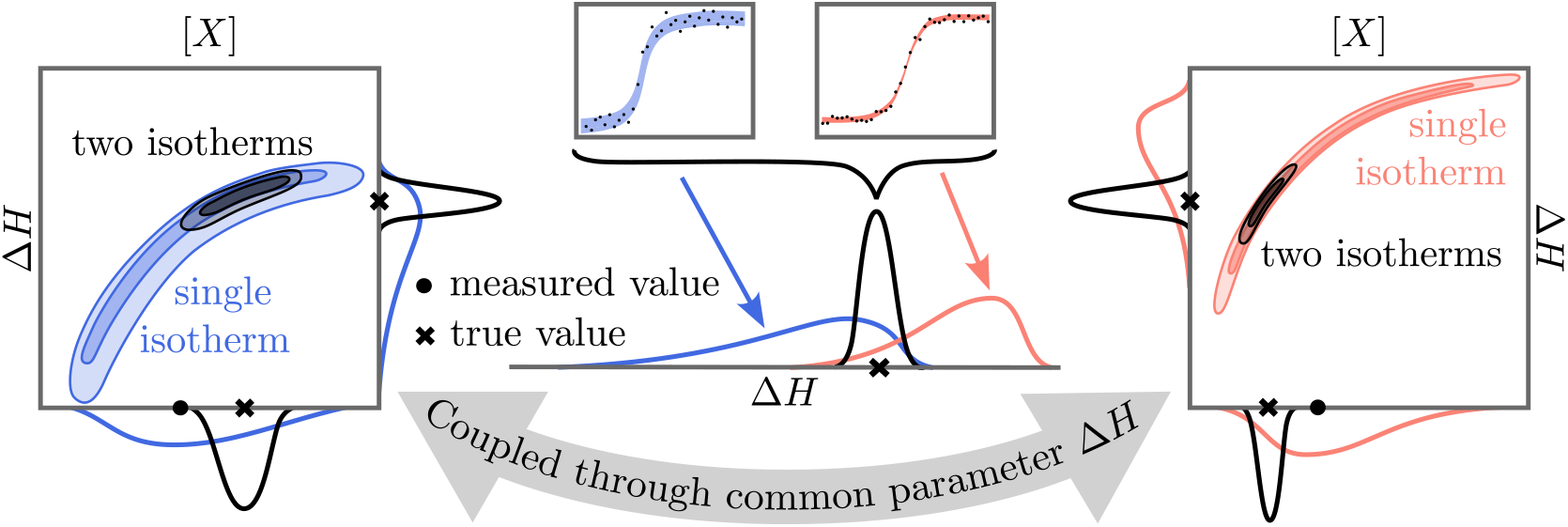
Resolving degeneracy via Bayesian analysis of multiple isotherms. Schematic posterior distributions for titrant concentration [*X*] and enthalpy Δ*H* based on single isotherms are indicated by the colored contours (far left and right). The posterior resulting from joint evaluation of two isotherms is shown in black. Since both isotherms are coupled via the enthalpy Δ*H*, a joint evaluation not only narrows down the respective thermodynamic parameters but also the analyte concentrations, e.g., [*X*].

#### 2.2.2 Parameter Counting

Generally, the parameters of primary interest are the thermodynamic binding parameters **Δ*G*** and **Δ*H***, which can be estimated by jointly analyzing multiple ITC datasets. In contrast, parameters such as the enthalpic offset Δ*H*_0_, analyte concentrations [***M***] and [***X***], and the measurement noise *σ* are considered dataset-specific nuisance parameters. When analyzing *N*_*I*_ independent datasets of the same system, the total number of model parameters becomes 2*N*_*S*_ + *N*_*I*_ (2 + *N*_*M*_ + *N*_*X*_). In the simple case of 1:1 binding (*N*_*S*_ = 1, *N*_*M*_ = *N*_*X*_ = 1), this equation reduces to 2 + 4*N*_*I*_. Thus, analyzing just one dataset requires the estimation of 6 parameters. In contrast, a more complex model such as dimeric binding analyzed across 9 datasets (as discussed in Sec. 3.4) involves estimating 58 parameters simultaneously.

As the dimensionality of the parameter space increases, parameter estimation using BI becomes significantly more challenging and resource intensive. The corresponding posterior distributions often exhibit strong correlations and complex geometries that hinder efficient sampling. High-dimensional spaces also make it more difficult to distinguish signal from noise and to ensure that the posterior is adequately explored. These challenges necessitate careful choices of priors, and the use of advanced sampling techniques to obtain robust and reliable estimates.

#### 2.2.3 Filtering of Concentration Ranges Based on Stoichiometry

In binding models with higher-order stoichiometries, it is sometimes necessary to impose additional constraints on the concentration priors to ensure that the inferred parameters reflect the correct binding stoichiometry. For instance, in an *s* : 1 binding model with the peptide in the syringe and the macro-molecule in the cell, we apply a filtering approach based on an estimate of the isotherm’s inflection point.

We approximate the inflection point by identifying the injection number *i* where the integrated heat Δ*Q*_*i*_ is closest to the midpoint value (Δ*H*_0_ + *Q*_1_)*/*2. Using this estimate, we compute the total concentrations of titrand and titrant in the cell after injection *i*, denoted as [*M*]_*t*_ and [*X*]_*t*_, respectively. To exclude solutions corresponding to lower stoichiometries, we enforce a constraint on their ratio by setting the prior probability to zero for any parameter set where [*X*]_*t*_*/*[*M*]_*t*_ *< s −* 0.5.

Although stoichiometry can also be restricted by placing tighter priors on Δ*G*_*j>*1_, we find that constraining the concentration priors is often more effective, especially for low stoichiometries, since this method has the advantage of not requiring prior estimates of the free energies. When analyzing multiple isotherms, it is often sufficient to apply this filtering to a subset of the data—particularly those isotherms where the inflection point is most clearly defined.

### 2.3 Three Bayesian Approaches for ITC Data

Accurately modeling uncertainty is essential for drawing reliable conclusions from ITC data. Experimental measurements inherently contain both random noise and systematic biases, which can affect not only the observed outputs—such as the integrated heats—but also critical experimental inputs, including analyte concentrations. BI provides a natural framework for incorporating and quantifying these uncertainties through the use of prior distributions, as introduced in Sec. 2.1.

To represent the absence of prior information, we selected broad uniform priors for the parameters of primary interest, **Δ*G*** and **Δ*H***, as well as for the nuisance parameters, Δ*H*_0_ and *σ*. The specific ranges used are provided in Tbl. 1.

**Table 1:** Priors chosen for model parameters.

| Parameter | Prior Type | Range |
| --- | --- | --- |
| $\Delta G_j$ (kcal/mol) | Uniform | $[-20 (b_j - 1), 0]$ |
| $\Delta H_j$ (kcal/mol) | Uniform | $[-20 (b_j - 1), 0]$ |
| $\Delta H_0$ (μcal) | Uniform | $[\Delta Q_1, 2\Delta Q_1 - \Delta Q_N]$ |
| $\sigma$ (μcal) | Uniform | $[0.001, 10]$ |
| $[M_l]$ (mM) | Lognormal | see Sec. 2.3 |
| $[X_k]$ (mM) | Lognormal | see Sec. 2.3 |

In contrast to these weakly informative priors, prior knowledge about analyte concentrations is often available in the form of laboratory measurements. A central challenge is how best to incorporate this information while accounting for measurement error and potential bias. Unlike uncertainties in the integrated heat, concentration uncertainties influence the control variables of the experiment and can propagate nontrivially into the inferred thermodynamic parameters.

Previous studies on calorimetry data have highlighted the significance of addressing measurement errors in concentrations [22, 25]. In particular, BI applied to single ITC datasets has been shown to effectively correct concentration estimates, provided that the analyte concentration prior is well modeled [22].

In this manuscript, we analyze three different approaches to modeling measurement uncertainties in analyte concentrations, aiming to establish a robust data analysis protocol for multiple isotherms. The three approaches are illustrated in Fig. 3 and described in the following.

**Figure 3:**
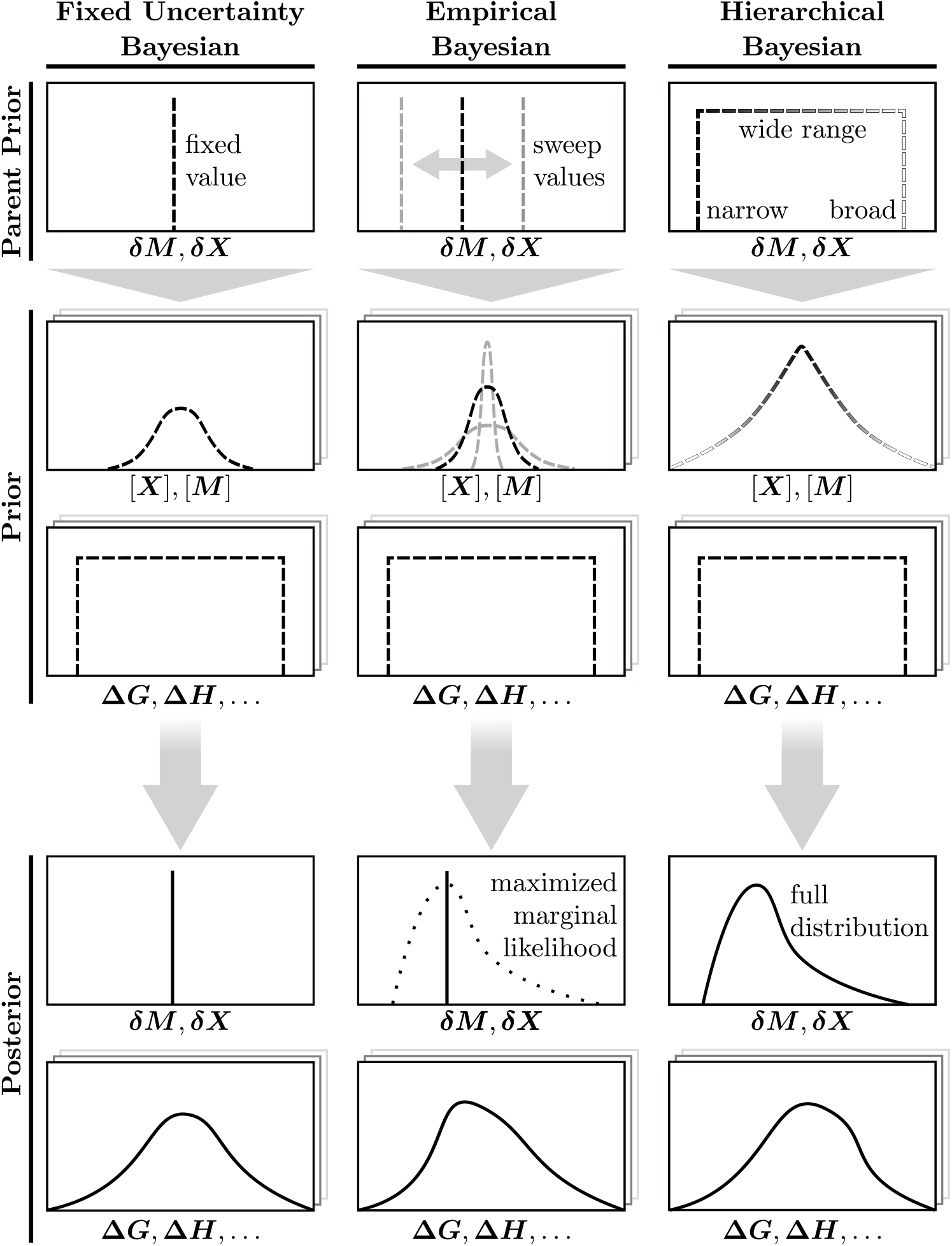
Three increasingly flexible strategies for modeling measurement uncertainties in analyte concentrations using BI. To model the prior for titrand and titrant concentrations, [***M***] and [***X***], all three approaches use lognormal priors centered on the measured analyte concentrations with widths determined by the relative uncertainty parameters ***δM*** and ***δX***, respectively. In the fixed uncertainty Bayesian approach, the uncertainties are specified in advance, based on prior experimental knowledge, and held constant across all datasets. The empirical Bayesian approach treats the uncertainty parameters as hyperparameters to be optimized by maximizing the marginal likelihood. The hierarchical Bayesian approach assigns prior distributions to the uncertainty parameters, which are then inferred along with all other model parameters, such as the thermodynamic parameters Δ*G* and Δ*H*.

#### 2.3.1 Fixed Uncertainty Bayesian

In the conventional fixed uncertainty Bayesian approach, concentration uncertainties for each analyte are assumed to be known and constant across all experiments. Specifically, the uncertainties ***δM*** and ***δX*** for the measured concentrations of titrant(s) and titrand(s) are fixed based on, *e*.*g*., prior experimental knowledge. These fixed values define independent lognormal priors for the analyte concentrations, centered at the measured values [***M***]_*m*_ and [***X***]_*m*_ with the fixed uncertainties ***δM*** and ***δX***.

This straightforward approach is computationally efficient and has been successfully applied in ITC parameter inference [22, 25, 11]. It is suitable for analyzing both individual ITC datasets and combined analyses of multiple datasets. However, if the true experimental variability differs from the assumed uncertainties, it may introduce bias in parameter estimates and error quantification (see Results, below).

#### 2.3.2 Empirical Bayesian

The empirical Bayesian approach, also known as maximum marginal likelihood, estimates concentration uncertainties directly from the data rather than assuming fixed values. Here, the uncertainty parameters ***δM*** and ***δX*** are treated as hyperparameters to be optimized. The posterior calculation is performed multiple times using different sets of hyperparameters and for the final parameter estimates the set of hyperparameters that maximizes the marginal likelihood is chosen. In practice, multiple posterior calculations can often be avoided by reweighting posterior distribution to reflect the differences in the priors, as introduced in Sec. 2.1.4.

This approach improves analyte concentration estimation by refining the lognormal distribution parameters based on observed variability. To achieve this, multiple datasets need to be analyzed in tandem to enable the estimation of concentration uncertainties. Compared to the fixed uncertainty method, this refinement increases robustness to variability in concentration errors. However, empirical Bayes shares limitations with traditional maximum likelihood methods, as it does not integrate over hyperparameter uncertainty, often resulting in overconfident parameter estimates (as seen below).

#### 2.3.3 Hierarchical Bayesian

The hierarchical Bayesian approach fully models concentration uncertainties as hyperparameters with their own prior distributions, enabling a probabilistic treatment of concentration variability across multiple datasets. In this framework, the uncertainty parameters ***δM*** and ***δX*** for each dataset are characterized by a “higher-level” distribution that is itself inferred from the data. The prior distribution is given by

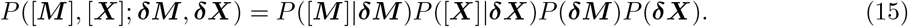

This approach naturally propagates the uncertainty in the analyte concentration errors through the inference process, resulting in more realistic credibility intervals (see Results, below), while also providing self-consistent feedback on the true measurement uncertainties. Computationally, the approach leads to increased complexity due to the need for sampling a higher-dimensional posterior distribution——specifically, *N*_*X*_ + *N*_*M*_ new variables corresponding to the concentration uncertainties of each analyte, assuming independent measurement errors. It also requires a choice for the prior distribution of the uncertainty parameters ***δM*** and ***δX***. In this manuscript, we employ a broad uniform distribution spanning measurement errors between 0% and 200%.

### 2.4 Synthetic Data Generation and Evaluation

To generate more realistic synthetic isotherms for testing our Bayesian framework, we incorporate measurement uncertainties in both heat observations and analyte concentrations. The observed heat is adjusted via

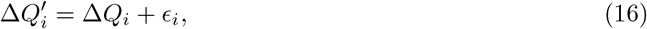

where *ϵ*_*i*_ represents measurement noise in Δ*Q*_*i*_, modeled as a normally distributed variable with zero mean and standard deviation *σ*. Additionally, uncertainty in analyte concentrations is introduced by sampling the synthetic measured titrand and titrant concentrations [***M***]_*m*_ and [***X***]_*m*_ from lognormal distributions centered at the ground truth values, with an associated uncertainty ***δM*** and ***δX***.

#### 2.4.1 1:1 Binding Model

For all synthetic isotherms generated using a 1:1 binding model, the ground-truth titrant concentration was fixed at [*X*] = 0.4 mM, while the titrand concentrations were distributed evenly between [*M*] = 0.02 mM and [*M*] = 0.06 mM, depending on the number of isotherms analyzed jointly. The synthetic measured values, [*M*]_*m*_ and [*X*]_*m*_, were shifted from the ground truth value using a lognormal distribution centered around the true values, with the spread determined by the measurement errors *δM* and *δX* in Tbl. 2. Furthermore, we set Δ*H*_0_ = 0 *µ*cal and the measurement noise was modeled with a standard deviation of *σ* = 1 *µ*cal. The synthetic injection protocol included an initial 2 *µ*l injection, which was excluded from the BI fit, followed by 35 injections of 10 *µ*l each at a temperature *T* = 25 °C.

**Table 2:** Synthetic parameter sets and Bayesian uncertainties. The table shows systems used to benchmark the different BI approaches to model experimental uncertainty. Each row lists input values for binding free energy Δ*G*, enthalpy Δ*H*, and analyte concentration measurement errors *δM* and *δX*, along with the resulting average 95% credible interval widths for Δ*G* and Δ*H* across 100 posterior samples obtained using the hierarchical model.

| System parameters |  |  |  | Bayesian findings |  |
| --- | --- | --- | --- | --- | --- |
| $\Delta G$<br>(kcal/mol) | $\Delta H$<br>(kcal/mol) | $\delta M$<br>(%) | $\delta X$<br>(%) | 95% $\Delta G$<br>(kcal/mol) | 95% $\Delta H$<br>(kcal/mol) |
| -9 | -11 | 10 | 10 | 0.23 | 2.9 |
| -9 | -11 | 20 | 20 | 0.32 | 5.3 |
| -9 | -11 | 20 | 5 | 0.23 | 2.9 |
| -9 | -15 | 10 | 10 | 0.19 | 3.8 |
| -9 | -7 | 10 | 10 | 0.32 | 1.9 |
| -11 | -11 | 10 | 10 | 0.89 | 2.8 |
| -7 | -11 | 10 | 10 | 0.30 | 3.6 |

The ranges of the prior distributions are denoted in Tbl. 1. The exact shape of the concentration priors are determined by the approach used to model the experimental uncertainty. The MCMC sampling was performed with the pocoMC package [32] using 4096 effective particles and eight times as many total independent samples. The initial runs were done employing the hierarchical approach, which was then reweighted to obtain the distributions for the other approaches, where of interest. General consistency among randomly performed independent MCMC replicas suggests that the posterior distributions are well sampled.

#### 2.4.2 1:2 Binding Model

For all synthetic isotherms generated using a 1:2 binding model, titrant concentrations were fixed at ground truth values of [*X*] = 0.6 mM for isotherms with dimer in the cell, and [*X*] = 0.2 mM for isotherms with dimer in the syringe. The titrand concentration was varied equally across [*M*] = 0.02, 0.04, 0.06 mM. In addition to the prior bounds specified in Tbl. 1, we imposed uniform priors on the coupling parameters ΔΔ*G* = Δ*G*_2_ −2Δ*G*_1_ ∈ [−5, 5] kcal*/*mol and ΔΔ*H* = Δ*H*_2_ −2Δ*H*_1_ ∈ [− 15, 15] kcal*/*mol. Furthermore, the concentration priors were constrained as described in Sec. 2.2.3 to enforce the correct binding stoichiometry during inference. All other simulation and run parameters were identical to Sec. 2.4.1. Importantly, rather than assigning separate uncertainties based on whether a species was in the cell or syringe, we assigned distinct uncertainties to the dimer and ligand themselves—reflecting that different binding partners may be subject to differing measurement precision, regardless of their experimental placement.

## 3 Results

We present results for Bayesian parameter inference applied to both synthetic and real ITC datasets. Synthetic data is used to compare different modeling approaches for experimental concentration uncertainties, assessing their validity, and evaluating parameter improvements when analyzing multiple datasets. Specifically, we examine the three approaches to modeling experimental concentration uncertainty outlined in Sec. 2.3, using synthetically generated ITC datasets for 1:1 binding. Additionally, we extend our analysis to alternative binding schemes, such as dimeric binding, which are discussed below. The hierarchical Bayesian approach is applied to true experimental data for both 1:1 and 2:1 stoichiometries.

### 3.1 Assessment of Bayesian Strategies for Concentration Measurement Uncertainty

To compare and validate the three approaches for modeling experimental concentration uncertainty, we evaluated 100 independent sets of three synthetic isotherms of a 1:1 binding model for various combinations of thermodynamic parameters and concentration measurement errors. Details of the parameter sets used can be found in Tbl. 2.

The validation plot in Fig. 4(b) shows the coverage of the true thermodynamic values as a function of their posterior percentiles. Ideally, a well-calibrated model would follow the diagonal, indicating that, for example, 95% credible intervals contain the true value 95% of the time. The fixed uncertainty approach exhibits a strong sensitivity to the assumed concentration measurement error used to construct the prior, leading to either under- or overconfident posteriors depending on the choice. The empirical Bayes method tends to produce mildly overconfident predictions, i.e., credibility intervals that cover the true value less often than their nominal percentiles. This may be because the approach fails to account for the uncertainty in the prior itself.

**Figure 4:**
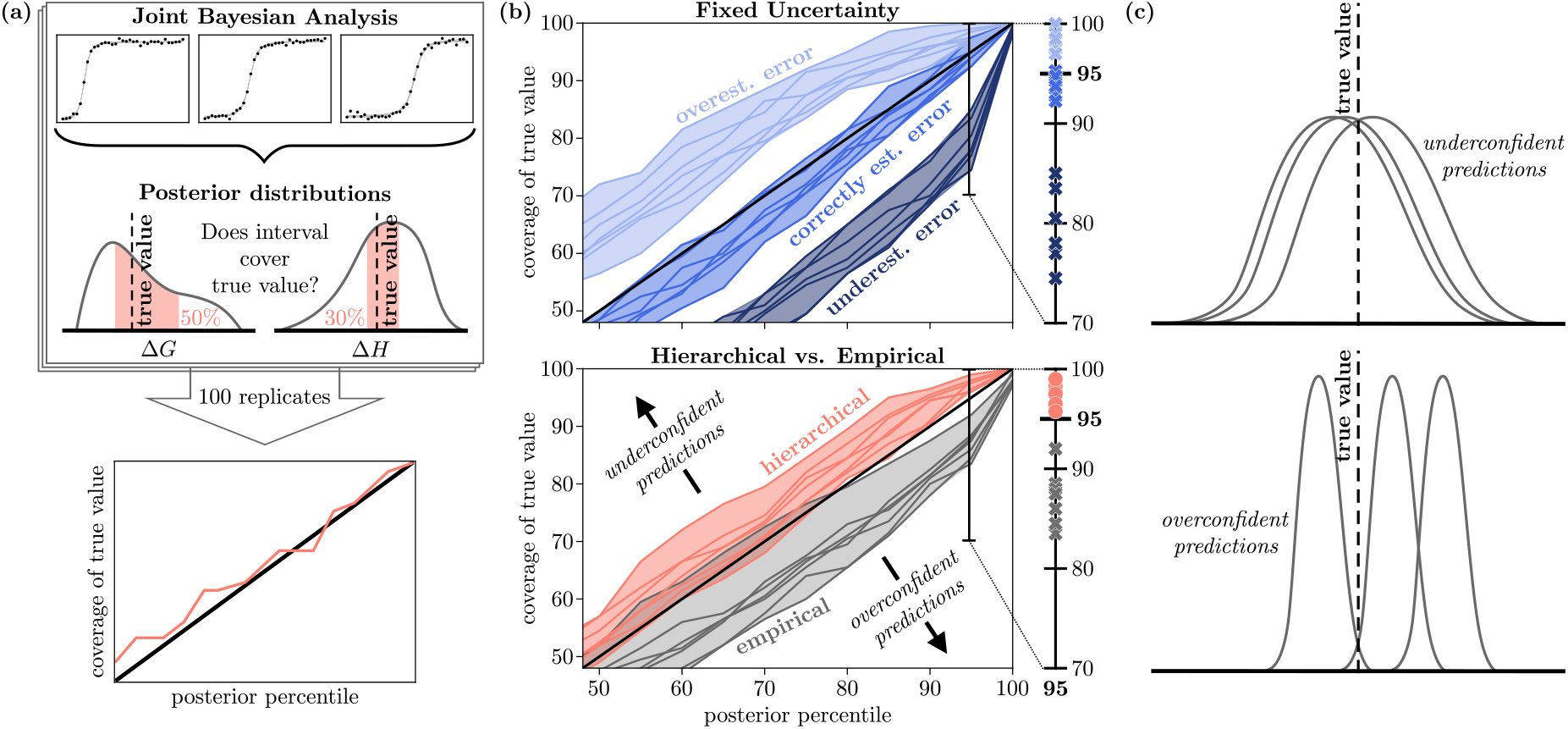
Comparison of three Bayesian approaches for modeling concentration uncertainty. Using synthetic data, the fixed-uncertainty, empirical, and hierarchical strategies are analyzed.**(a)** Schematic of the validation pipeline used to generate the plots. **(b)** Validation plots comparing the three Bayesian approaches, with ideal behavior conforming to the diagonal. Each line represents an average over 100 independent BI runs for one set of system parameters listed in Tbl. 2, each analyzing three independent isotherms. For the fixed-uncertainty method, we compare three scenarios: accurate uncertainty estimation, overestimation (double the true uncertainty), and underestimation (half the true uncertainty). **(c)** Conceptual illustrations of underconfident and overconfident predictions.

The hierarchical model yields slightly conservative predictions, i.e., “underconfident” intervals which are slightly broader than the actual behavior. Given that perfect accuracy in uncertainty estimation is not possible with finite data, having slightly underconfident intervals seems ideal. The broadened intervals may reflect the added uncertainty arising from estimating prior hyperparameters across datasets.

For the subsequent analysis, we adopt the hierarchical approach due to its principled treatment of prior uncertainty and its consistent performance across varying experimental conditions. Although its credible intervals are slightly conservative, this behavior is preferable in the context of ITC modeling, where overconfident estimates may obscure true uncertainty and result in misleading interpretations, see Fig. 4(c).

Table 2 contains the average 95% credible interval widths for Δ*G* and Δ*H* based on the hierarchichal evaluation. While there is some variation across conditions, the credible interval widths generally increase with larger concentration uncertainties, highlighting the strong influence of concentration uncertainty on inference quality.

### 3.2 Analyzing Multiple Datasets

To assess how parameter inference improves with increasing dataset size, we conducted 100 independent hierarchical BI runs using synthetic sets of *n* = 2, 3, 4, …, 9 isotherms for the first parameter set in Tbl. 2. We compared the resulting posterior distributions in terms of both accuracy and uncertainty. In addition, we highlight the qualitative differences between single-isotherm analyses and a joint analysis of multiple isotherms.

Figure 5(a) compares the one-dimensional posterior marginals for free energy and enthalpy obtained from independent single-dataset runs with those from a joint hierarchical Bayesian fit. Since the hierarchical model infers concentration measurement uncertainty directly from the data, as illustrated in Fig. 6(a), its benefits only manifest when multiple isotherms are jointly analyzed. To ensure a fair comparison, the single-dataset runs were performed using a fixed uncertainty prior that matched the synthetic measurement error of 10% used to generate the data.

**Figure 5:**
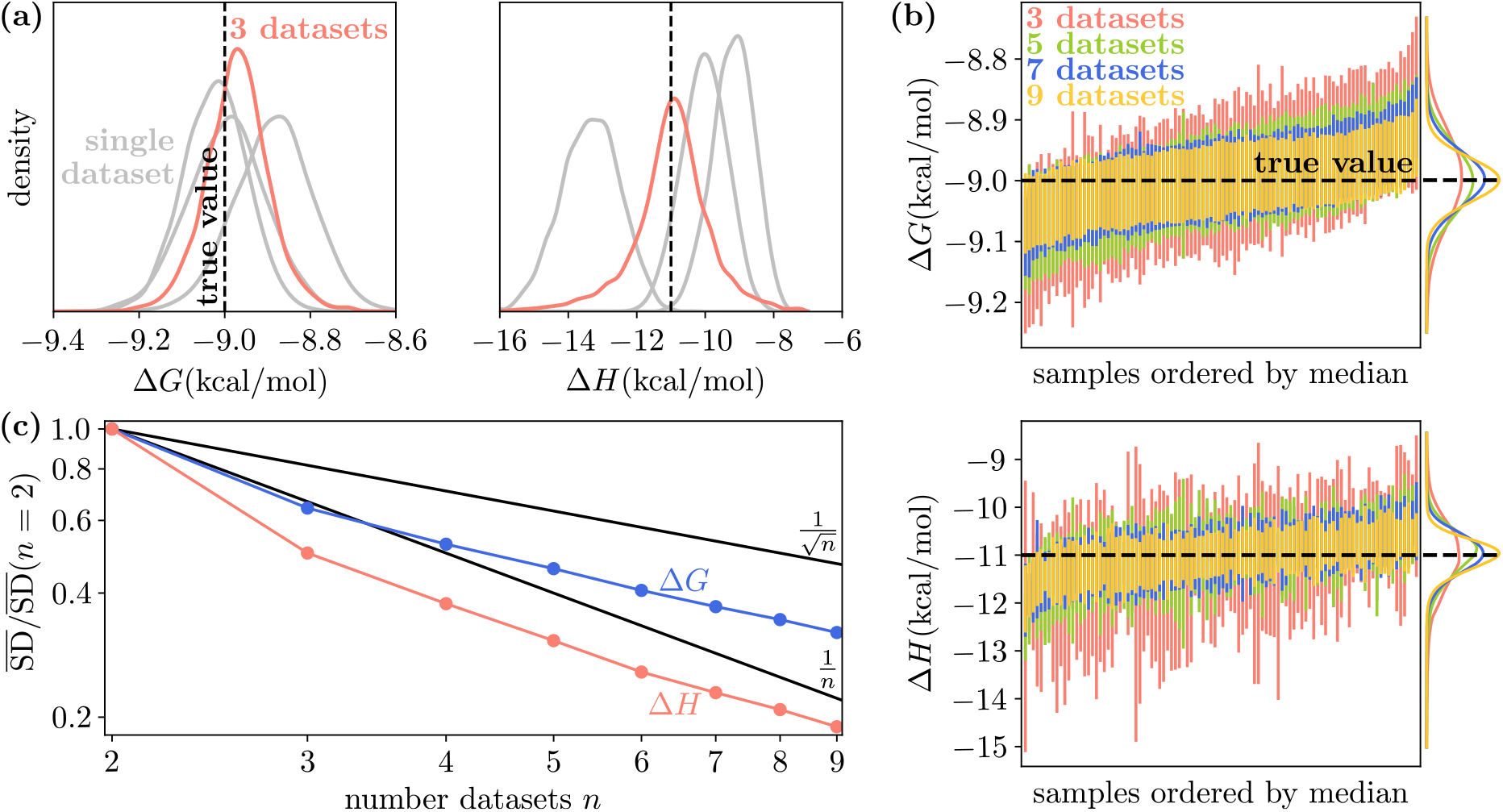
Improvement of posterior precision increasing dataset size. Analysis is performed on synthetic isotherms generated with Δ*G* = −9 kcal/mol, Δ*H* = −11 kcal/mol, and relative concentration uncertainties *δM* = *δX* = 10%. **(a)** Example posterior marginal distributions from single-dataset inferences compared to the result from a joint hierarchical Bayesian analysis. **(b)** 95% credible intervals for Δ*G* and Δ*H* inferred from hierarchical fits using 3, 5, 7, and 9 datasets, for 100 independent synthetic replicates. **(c)** Mean posterior standard deviation for Δ*G* and Δ*H* as a function of the number of datasets used in the hierarchical model. The uncertainty in Δ*H* decreases faster than the central limit 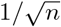 scaling, reflecting an accelerated gain in precision due to hierarchical information sharing.

**Figure 6:**
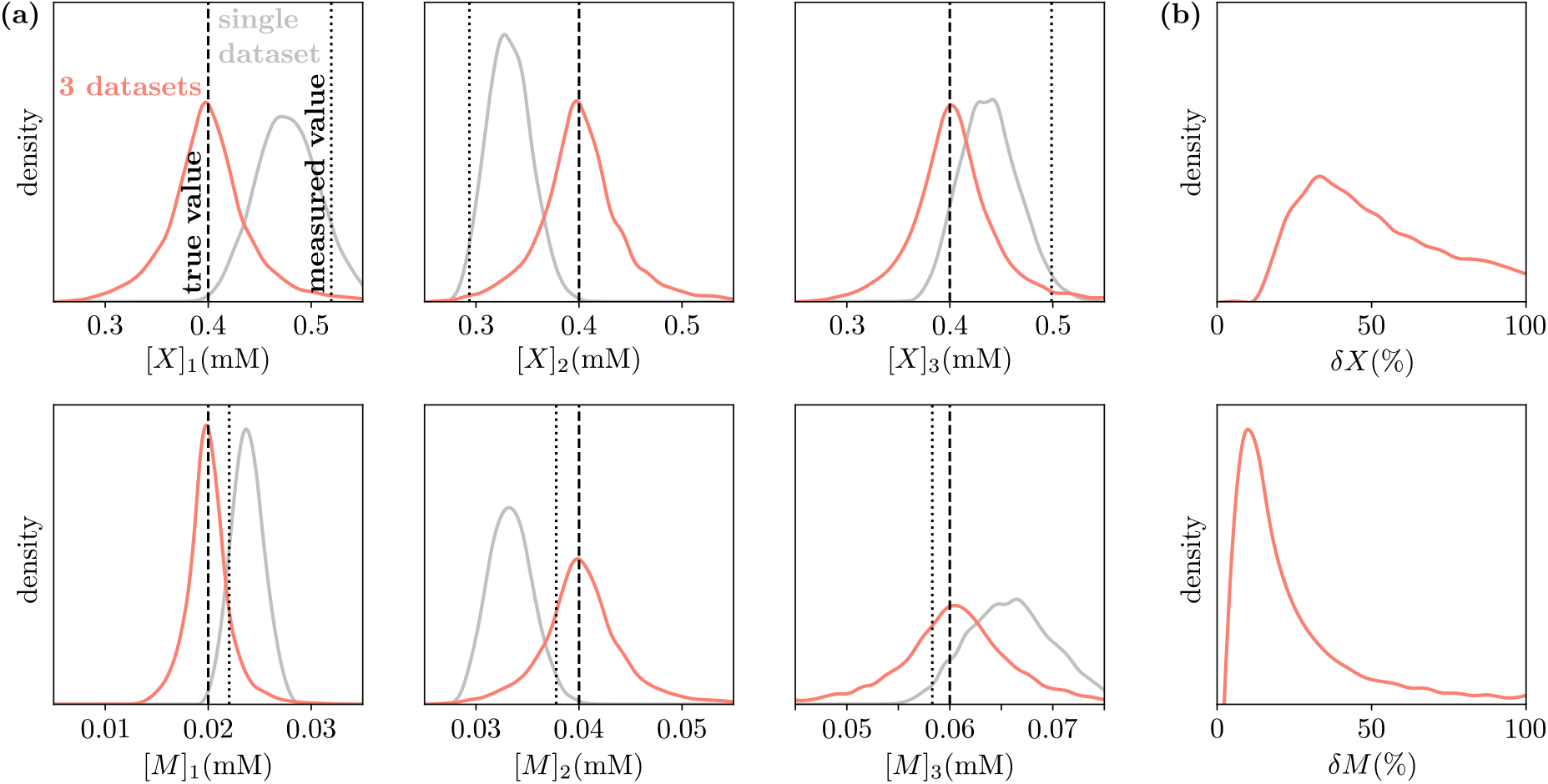
Example of posterior distributions for analyte concentrations and their measurement uncertainties based on synthetic isotherms generated with Δ*G* = −9 kcal/mol, Δ*H* = −11 kcal/mol, and relative uncertainties *δM* = *δX* = 10%. **(a)** Posterior distributions for titrant and titrand concentrations [*X*] and [*M*] from single-dataset Bayesian fits as well as a joint hierarchical Bayesian fit. **(b)** Posterior distributions for the measurement uncertainties in titrant and titrand concentrations, *δX* and *δM*, respectively. The larger posterior values of *δX* reflect the greater-than-expected measurement errors for titrant concentrations that are nevertheless fitted accurately by the hierarchical approach.

The single-dataset results show consistent estimates for free energy across runs but exhibit substantial variation in the enthalpy and concentration marginals. On the other hand, the hierarchical analysis leads to a modest improvement in free energy estimation but delivers significantly more accurate and concentrated posteriors for enthalpy and analyte concentrations. This behavior is consistent with the degeneracy relationship described in Eq. (13) and illustrated in Fig. 2, where enthalpy and analyte concentrations scale linearly with the degeneracy parameter *α*, while free energy is affected only logarithmically. These results demonstrate that parameters more susceptible to the effects of the degeneracy—such as enthalpy and concentration—derive the greatest benefit from multi-isotherm analysis, where shared information across datasets helps constrain otherwise weakly identifiable quantities.

In addition, the posterior distributions for the measurement uncertainties in Fig. 6(b) highlight how the hierarchical approach enables direct inference of the true analyte concentration variation across experiments. Here, the larger posterior values of *δM* reflect the greater-than-expected measurement errors for titrant concentrations that are nevertheless fitted accurately by the hierarchical approach.

To further investigate the influence of dataset number on posterior uncertainty, we performed additional hierarchical Bayesian analyses using *n* = 2, 3, 4, …, 9 isotherms for Δ*G* = −9 kcal/mol, Δ*H* = −11 kcal/mol, and *δM* = *δX* = 10 %. Figure 5(b) shows the resulting 95% credible intervals from the three-, five-, seven-, and nine-isotherm runs. As expected, incorporating additional datasets leads to a systematic reduction in posterior uncertainty, yielding increasingly precise parameter estimates. Notably, the inclusion of more isotherms enhances the hierarchical model’s ability to infer measurement uncertainty, which in turn mitigates the higher variability observed in enthalpy estimates under smaller dataset sizes. This is further highlighted in Fig. 5(c) which shows the mean posterior standard deviation for Δ*G* and Δ*H* as a function of dataset count. The uncertainty in Δ*H* decreases more rapidly than the canonical 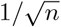 central limit scaling, particularly at small *n*, indicating a pronounced precision gain from hierarchical information sharing. In contrast, the standard deviation of Δ*G* approaches the central limit behavior more quickly, suggesting that fewer datasets are required for stable estimation of binding free energy.

### 3.3 Mg(II) - EDTA Binding

Following the validation with synthetic data, we applied the hierarchical Bayesian approach to experimental ITC data. Our analysis is based on a publicly available dataset originally published in Ref. [22], which investigates the 1:1 binding interaction between Mg(II) and the chelator EDTA. To rigorously characterize experimental uncertainty, the original study repeated the entire ITC experiment 14 times, with each replicate involving independently prepared solutions of titrant (MgCl_2_), titrand (EDTA), and buffer (50 mM Tris-HCl, pH 8.0). This careful design ensured that any concentration errors were uncorrelated across replicates. The first nine replicates used titrant and titrand concentrations of [*X*] = 0.5 mM and [*M*] = 0.05 mM, respectively, while the remaining five used [*X*] = 1.0 mM and [*M*] = 0.1 mM.

As a baseline, we analyzed each of the 14 isotherms individually using a fixed uncertainty prior, assuming a 10% measurement error as previously proposed in Ref. [22]. We then performed combined hierarchical analyses by randomly grouping the isotherms into sets of three, five, and seven. Finally, we carried out a joint analysis using all 14 datasets.

The results for Mg(II)-EDTA binding are consistent with findings for synthetic data. Fig. 7 shows that the full hierarchical analysis of 14 datasets produces markedly more precise estimates, with Δ*G* = −8.99(4) kcal/mol and Δ*H* = −2.19(8) kcal/mol, respectively, as compared to using fewer datasets. Posteriors for 3, 5, and 7 datasets show the expected monotonic increase in precision, with mutual agreement among the distributions likewise increasing. The posterior distributions for the measurement uncertainties are shown in Fig. 7(c), yielding *δX* = 9(4) % and *δM* = 12(5) %, where the numbers in parentheses indicate the 95% credible interval half-widths, highlighting that the originally assumed 10% error was indeed a reasonable approximation.

**Figure 7:**
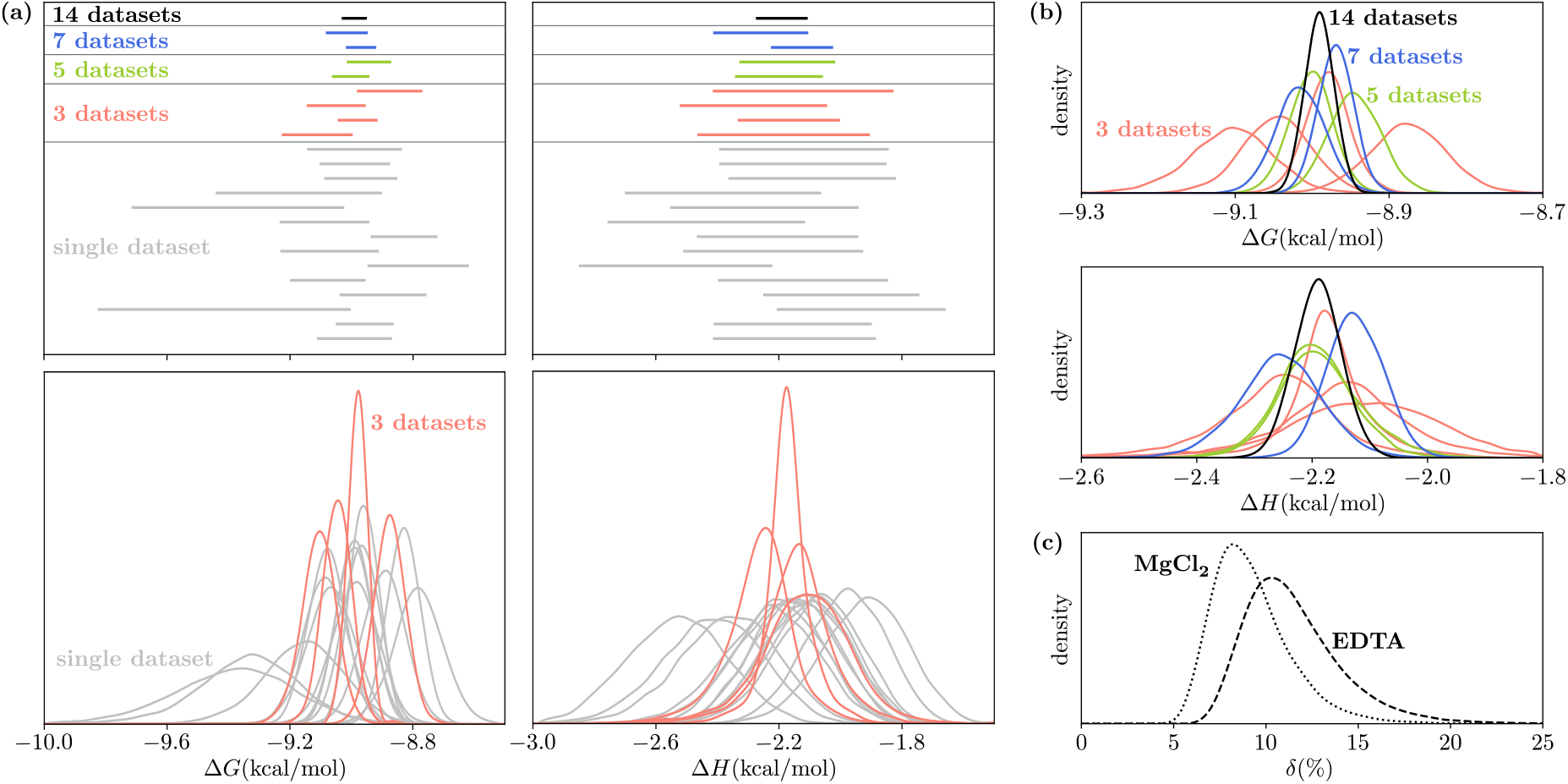
Analysis of 14 isotherms of the 1:1 binding interaction between Mg(II) and the chelator EDTA using hierarchichal BI. **(a**,**b)** 95% credible intervals and full marginal posterior distributions of thermodynamic parameters based on the combined hierarchical Bayesian analyses of randomly grouped sets of three, five, and seven isotherms as well as a full hierarchical run using all 14 isotherms and separate single-dataset analyses. **(c)** Posterior distributions for the measurement uncertainties of titrand and titrant concentrations based on the full hierarchical Bayesian analysis.

### 3.4 Dimeric Binding

To evaluate the performance of the hierarchical Bayesian model for more complex binding schemes and to assess the variability in parameter estimation across different conditions, we analyzed 108 independent sets of three synthetic isotherms generated under a 2:1 binding model. Thermodynamic parameters of the mono-ligated species were set to Δ*G*_1_ = *−*9 kcal*/*mol and Δ*H*_1_ = *−*11 kcal*/*mol, with concentration uncertainties fixed at 10%. The cooperativity parameters ΔΔ*G* = Δ*G*_2_ *−* 2Δ*G*_1_ and ΔΔ*H* = Δ*H*_2_ −2Δ*H*_1_, describing the difference between the first and second binding event, were varied across nine distinct parameter sets as detailed in Tbl. 3. Specifically, we investigated how the placement of the dimer—either in the syringe or in the cell—affects parameter estimation. For each thermodynamic parameter set, the 108 isotherms were, thus, evenly divided among sets featuring the dimer in the cell 3, 2, 1, or 0 times.

**Table 3:**
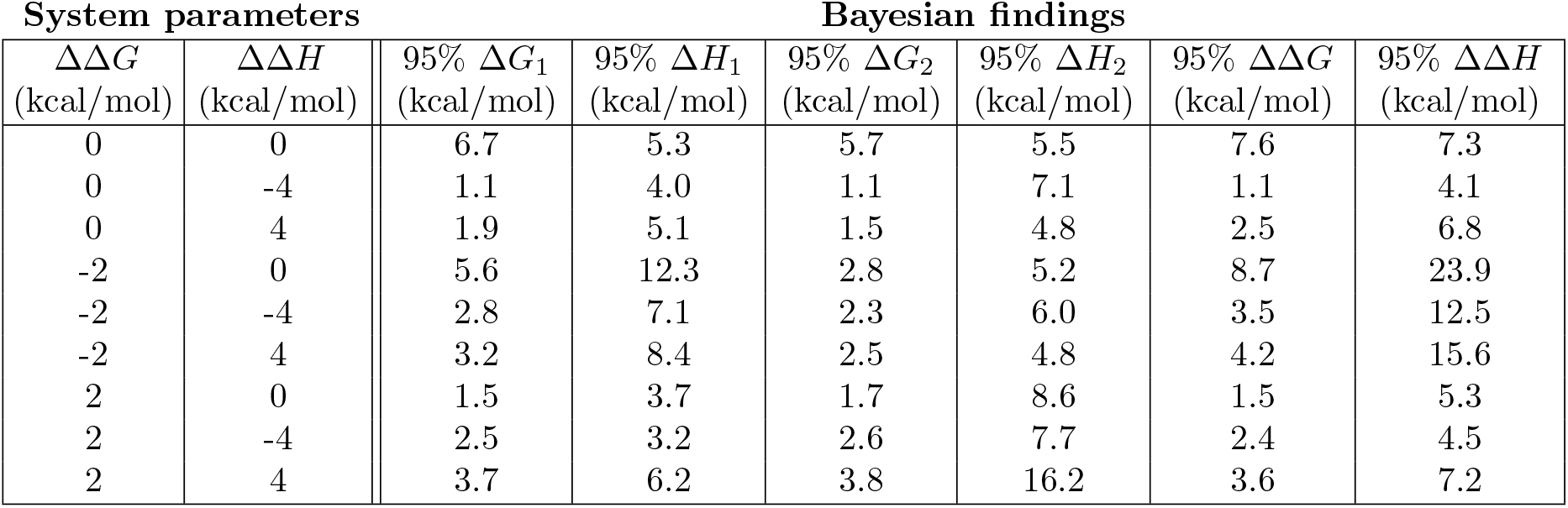
Synthetic parameter sets and Bayesian uncertainties. The table shows systems used to analyze the variability in parameter estimation across different conditions for a 2:1 binding scheme using synthetic data with Δ*G*_1_ = −9 kcal/mol, Δ*H*_1_ = −11 kcal/mol, and analyte concentration measurement errors *δM* = *δX* = 10 %. Each row lists input values for the thermodynamic cooperativity parameters ΔΔ*G* and ΔΔ*H*, along with the resulting average 95% credible interval widths for Δ*G*_*j*=1,2_, Δ*H*_*j*=1,2_ as well as ΔΔ*G* and ΔΔ*H* across 108 posterior samples obtained using the hierarchical model.

| System parameters |  | Bayesian findings |  |  |  |  |  |
| --- | --- | --- | --- | --- | --- | --- | --- |
| $\Delta\Delta G$<br>(kcal/mol) | $\Delta\Delta H$<br>(kcal/mol) | 95% $\Delta G_1$<br>(kcal/mol) | 95% $\Delta H_1$<br>(kcal/mol) | 95% $\Delta G_2$<br>(kcal/mol) | 95% $\Delta H_2$<br>(kcal/mol) | 95% $\Delta\Delta G$<br>(kcal/mol) | 95% $\Delta\Delta H$<br>(kcal/mol) |
| 0 | 0 | 6.7 | 5.3 | 5.7 | 5.5 | 7.6 | 7.3 |
| 0 | -4 | 1.1 | 4.0 | 1.1 | 7.1 | 1.1 | 4.1 |
| 0 | 4 | 1.9 | 5.1 | 1.5 | 4.8 | 2.5 | 6.8 |
| -2 | 0 | 5.6 | 12.3 | 2.8 | 5.2 | 8.7 | 23.9 |
| -2 | -4 | 2.8 | 7.1 | 2.3 | 6.0 | 3.5 | 12.5 |
| -2 | 4 | 3.2 | 8.4 | 2.5 | 4.8 | 4.2 | 15.6 |
| 2 | 0 | 1.5 | 3.7 | 1.7 | 8.6 | 1.5 | 5.3 |
| 2 | -4 | 2.5 | 3.2 | 2.6 | 7.7 | 2.4 | 4.5 |
| 2 | 4 | 3.7 | 6.2 | 3.8 | 16.2 | 3.6 | 7.2 |

Table 3 reveals substantial variability in the precision of thermodynamic parameter estimates across different cooperativity scenarios, underscoring the challenges of inferring coupled binding behavior. For example, the case ΔΔ*G* = 0 kcal/mol, ΔΔ*H* = −4 kcal/mol estimates the coupling parameters ΔΔ*G* and ΔΔ*H* with uncertainties of 1.1 and 4.1 kcal/mol, respectively—roughly 8-fold and 6-fold smaller than the corresponding uncertainties in the case ΔΔ*G* = −2 kcal/mol, ΔΔ*H* = 0 kcal/mol. In general, a negative ΔΔ*G* seems to hinder the accurate estimation of ΔΔ*H*.

Compared to the 1:1 binding model results in Tbl. 2, the 2:1 model yields broader credible intervals overall. Notably, parameter uncertainty is often greater for the first binding event than for the second. For instance, in the previously noted case ΔΔ*G* = *−*2 kcal/mol, ΔΔ*H* = 0 kcal/mol, the 95% credible intervals for Δ*G*_1_ and Δ*H*_1_ are 5.6 and 12.3 kcal/mol, respectively—substantially wider than those for Δ*G*_2_ (2.8 kcal/mol) and Δ*H*_2_ (5.2 kcal/mol). Moreover, since the second binding event corresponds to larger absolute thermodynamic values, its relative uncertainty is often markedly lower. This trend highlights that the second binding step tends to be more precisely identifiable than the intermediate step, both in free energy and enthalpy.

To illustrate the dependence of the parameter estimation on the position of the dimer in the cell or the syringe, Fig. S1 displays the accumulated posterior marginals across parameter sets featuring the dimer in the cell 3, 2, 1, or 0 times. The most narrow fits of the thermodynaic parameters are achieved for the sets featuring the dimer in the cell three times, while the sets featuring the dimer consistently in the syringe, lead too considerably wider distributions.

### 3.5 LC8 and Multiple Binding Partners

To demonstrate the versatility of our Bayesian pipeline, we applied the hierarchical Bayesian approach—along with the insights developed in Sec. 3.4—to experimental dimeric ITC data. We examined binding of the hub protein LC8 to client peptides generated from endogenous proteins Swallow and BIM, as well as VP35 from Ebola virus.

The datasets for BIM and Swallow, recorded at 10 °C, have been previously published in Refs. [37, 38], and consist of 9 and 8 isotherms, respectively. Experiments were conducted in a buffer containing 50 mM sodium phosphate, 50 mM sodium citrate, 50 mM sodium chloride, and 1 mM sodium azide (pH 5.5). The VP35 dataset, recorded at 25 °C and previously published in Ref. [39], comprises 4 isotherms. The corresponding buffer solution contained 50 mM sodium phosphate, 50 mM sodium chloride, 1 mM sodium azide, and 5 mM *β*-mercaptoethanol (pH 7.5).

Titrant and titrand concentrations, as well as the isotherms and their fits based on the posterior distributions, are shown in Fig. S2. Each set of isotherms was analyzed using the hierarchical Bayesian pipeline described in Sec. 2.4.2. The posterior estimates for the thermodynamic parameters are shown in Tbl. 4 and visualized in Fig. 8.

**Table 4:**
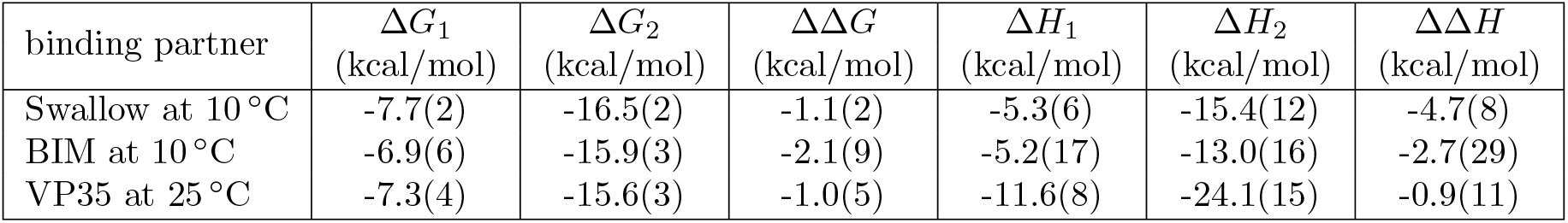
Posterior estimates and 95% credible interval half-widths for the thermodynamic binding parameters of LC8 and its three binding partners Swallow at 10 °C, BIM at 10 °C, and VP35 at 25 °C.

| binding partner | $\Delta G_1$<br>(kcal/mol) | $\Delta G_2$<br>(kcal/mol) | $\Delta\Delta G$<br>(kcal/mol) | $\Delta H_1$<br>(kcal/mol) | $\Delta H_2$<br>(kcal/mol) | $\Delta\Delta H$<br>(kcal/mol) |
| --- | --- | --- | --- | --- | --- | --- |
| Swallow at 10 °C | -7.7(2) | -16.5(2) | -1.1(2) | -5.3(6) | -15.4(12) | -4.7(8) |
| BIM at 10 °C | -6.9(6) | -15.9(3) | -2.1(9) | -5.2(17) | -13.0(16) | -2.7(29) |
| VP35 at 25 °C | -7.3(4) | -15.6(3) | -1.0(5) | -11.6(8) | -24.1(15) | -0.9(11) |

**Figure 8:**
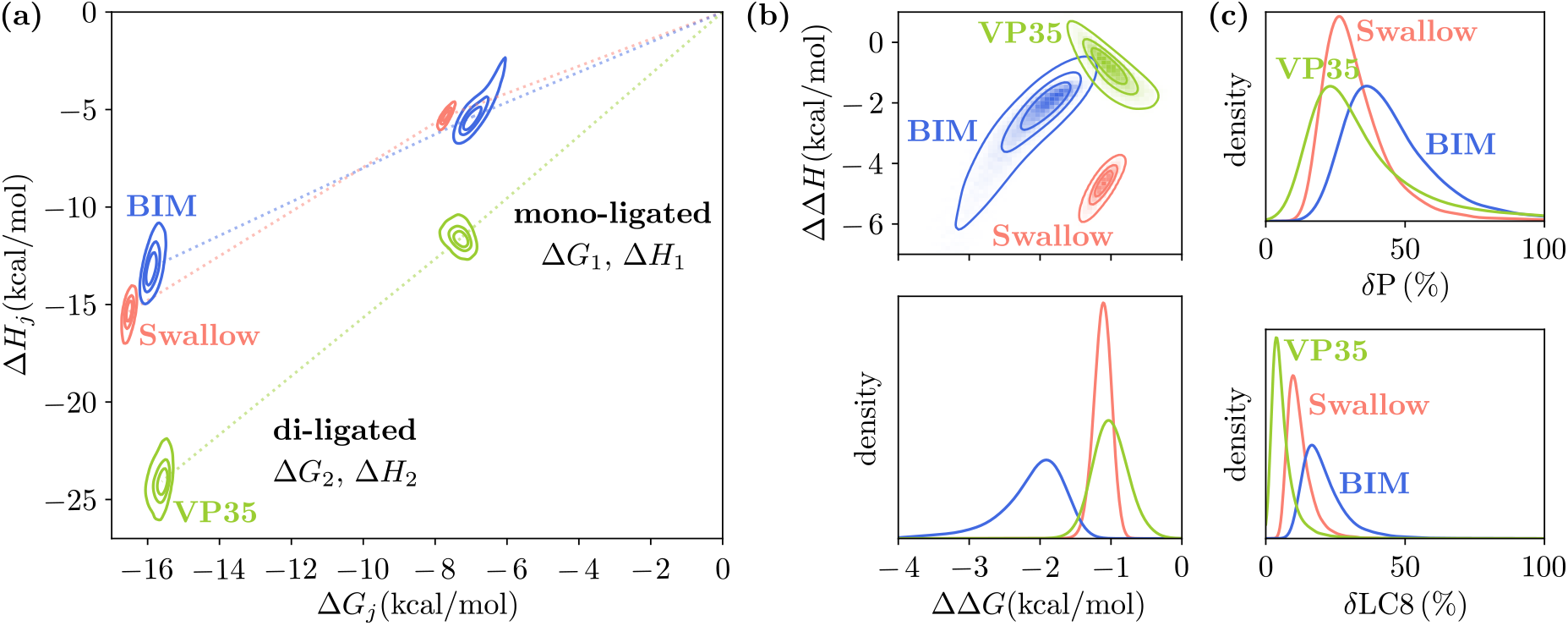
Quantitative analysis of cooperative dimeric binding. Hierarchical Bayesian inference was performed on ITC data obtained for the hub protein LC8 and its binding partners: Swallow at 10 °C, BIM at 10 °C, and VP35 at 25 °C. **(a)** Posterior distributions of the free energies Δ*G*_*j*_ and enthalpy Δ*H*_*j*_ for the mono-ligated (*j* = 1) and di-ligated (*j* = 2) states. Contour lines enclose the 39^th^, 68^th^, and 95^th^ percentiles of the joint probability density. **(b)** Joint and marginal posterior distributions of the coupling free energy ΔΔ*G* = Δ*G*_2_ *−* 2Δ*G*_1_ and coupling enthalpy ΔΔ*H* = Δ*H*_2_ *−* 2Δ*H*_1_, quantifying the cooperativity of the second binding event. The distributions reveal ligand-specific differences in binding thermodynamics and cooperativity. **(c)** Posterior distributions for the measurement uncertainties of the binding partners (*δ*P) and LC8 (*δ*LC8).

The results for all three binding partners indicate cooperative binding at the second site, but also reveal distinct, ligand-specific behaviors. BIM exhibits the strongest cooperative affinity, exceeding that of Swallow and VP35, whereas VP35 shows significantly larger total binding enthalpies for both the first and second binding events. Notably, the precision of parameter estimates does not depend solely on the number of isotherms or on whether the dimer was placed in the cell or syringe: the four isotherms used for VP35 yielded results comparable in precision to those of BIM, which relied on nine isotherms. Additionally, Fig. 8(c) illustrates a broad range of concentration measurement errors across experiments, with mean estimated uncertainties for LC8 and its partners ranging from 6% to 46%.

## 4 Discussion

Obtaining robust and interpretable estimates of parameters and their uncertainties remains a fundamental challenge in the analysis of biophysical data, particularly in the face of noisy measurements or uncertain experimental conditions. Isothermal titration calorimetry (ITC) exemplifies these challenges: the inference of thermodynamic parameters such as binding affinity and enthalpy is complicated by strong coupling between thermodynamic parameters and analyte concentrations, which are often only approximately known. In this study, we demonstrate that hierarchical Bayesian inference (BI) offers a powerful and flexible framework for addressing these issues. By jointly analyzing replicate isotherms and inferring both concentration values and their uncertainties directly from the data, hierarchical BI consistently outperforms approaches that rely on fixed or empirically corrected concentrations, yielding more accurate and reliable posterior distributions.

Hierarchical BI can largely overcome a key limitation of Bayesian inference, namely the need to specify prior distributions for parameters. By placing additional priors over analyte concentration uncertainties, the hierarchical method removes the need for precise independent estimates of measurement error—an aspect that can otherwise lead to significant under- or overestimation of uncertainty in thermodynamic parameters. Moreover, the hierarchical approach enables direct inference of the true analyte concentrations and their variation across experiments, which can be valuable even when nominal concentrations are reported with well known precision as small datasets can yield a wide range of effective variance.

It should be noted that the hierarchical approach performs best when analyte concentrations are prepared independently across isotherms, as this allows the model to infer individual latent concentrations constrained by a shared population-level prior. When the same solution is reused across experiments, this assumption no longer holds, and the model must be adapted to account for shared or correlated concentration values, for example by assigning a common latent variable to linked experiments.

An important strength of our Bayesian pipeline is its expandability to complex binding scenarios. Cooperative, competitive, and multivalent binding models can be analyzed by the software we have developed [33].

Despite the strengths of hierarchical BI, its computational demands remain a significant consideration. After evaluating multiple Monte Carlo sampling strategies, we selected the sophisticated and highly parallelized engine implemented in the pocoMC package [32] for our analysis. Nonetheless, resource requirements scale steeply with both the number of isotherms and the complexity of the binding model, which can limit scalability and highlights the need to balance model complexity, dataset size, and available computational resources. For instance, analyzing three isotherms of a 1:1 binding model required approximately 90 minutes on a four-core system, whereas analyzing seven isotherms under identical sampling settings extended the runtime to nearly 11 hours.

In future work, we plan to continue our exploration of methods for estimating posteriors for multiple datasets based on combining MCMC runs for data subsets [27].

The potential value of analyzing synthetic data for designing experiments is important to note. As models grow in complexity, more datasets are generally required to distinguish between competing hypotheses and constrain correlated parameters. For example, we show that simulating different titration setups—such as placing the dimer in the syringe versus in the cell for a 2:1 binding model—can provide practical insight into which experiments or which combination of experiments are most informative. BI can serve not only as an analysis method but also as a framework for experimental planning for complex scenarios.

Although this manuscript is focused on ITC, the methods and principles demonstrated here extend naturally to other areas of biophysics. Hierarchical BI is particularly valuable for any replicated experimental system in which key parameters are influenced by poorly constrained or variable experimental conditions—such as transporter kinetics, enzyme turnover assays, or systems biology models. In such cases, nuisance parameters like substrate concentrations, protein copy numbers, or measurement baselines may introduce significant uncertainty and correlation. As shown in previous work on biochemical networks and signaling pathways [20, 21], Bayesian approaches offer powerful tools for handling such complexity. Our findings support and extend this perspective, emphasizing how hierarchical modeling enables principled, data-driven learning of both parameters of interest and the underlying experimental uncertainties that shape them.

## 5 Conclusion

In this work, we have demonstrated that hierarchical Bayesian inference (BI) provides a robust and flexible framework for extracting thermodynamic parameters from isothermal titration calorimetry (ITC) data. By jointly modeling replicate isotherms and inferring latent concentrations and their uncertainties alongside binding parameters, hierarchical BI surpasses traditional approaches that rely on nonlinear least squares fitting or fixed concentrations and uncertainties, yielding more accurate and interpretable posterior estimates.

Our results highlight the advantages of hierarchical modeling in reducing reliance on precise prior knowledge of concentration uncertainties, and enabling direct inference of concentration variability. While computational demands remain a practical consideration—especially for complex binding models and larger datasets—the benefits in terms of improved parameter identifiability and uncertainty quantification are large. Furthermore, the flexibility of the Bayesian framework facilitates its extension to more complex binding schemes and supports the design of more informative experiments through simulation.

Importantly, although this study focuses on ITC, the methodological insights extend broadly to other biophysical and biochemical systems characterized by poorly constrained or variable experimental conditions that influence key parameters of interest. Hierarchical Bayesian modeling thus represents a powerful method for integrative data analysis and experimental planning, fostering more reliable and nuanced interpretations of complex biological measurements.

## Acknowledgments

The authors are grateful for support from the National Institutes of Health under Grant GM141733 to E.J.B. and D.M.Z and from the National Science Foundation under Grant MCB 2119837 to D.M.Z.

## Supplemental Figures

**Figure S1:**
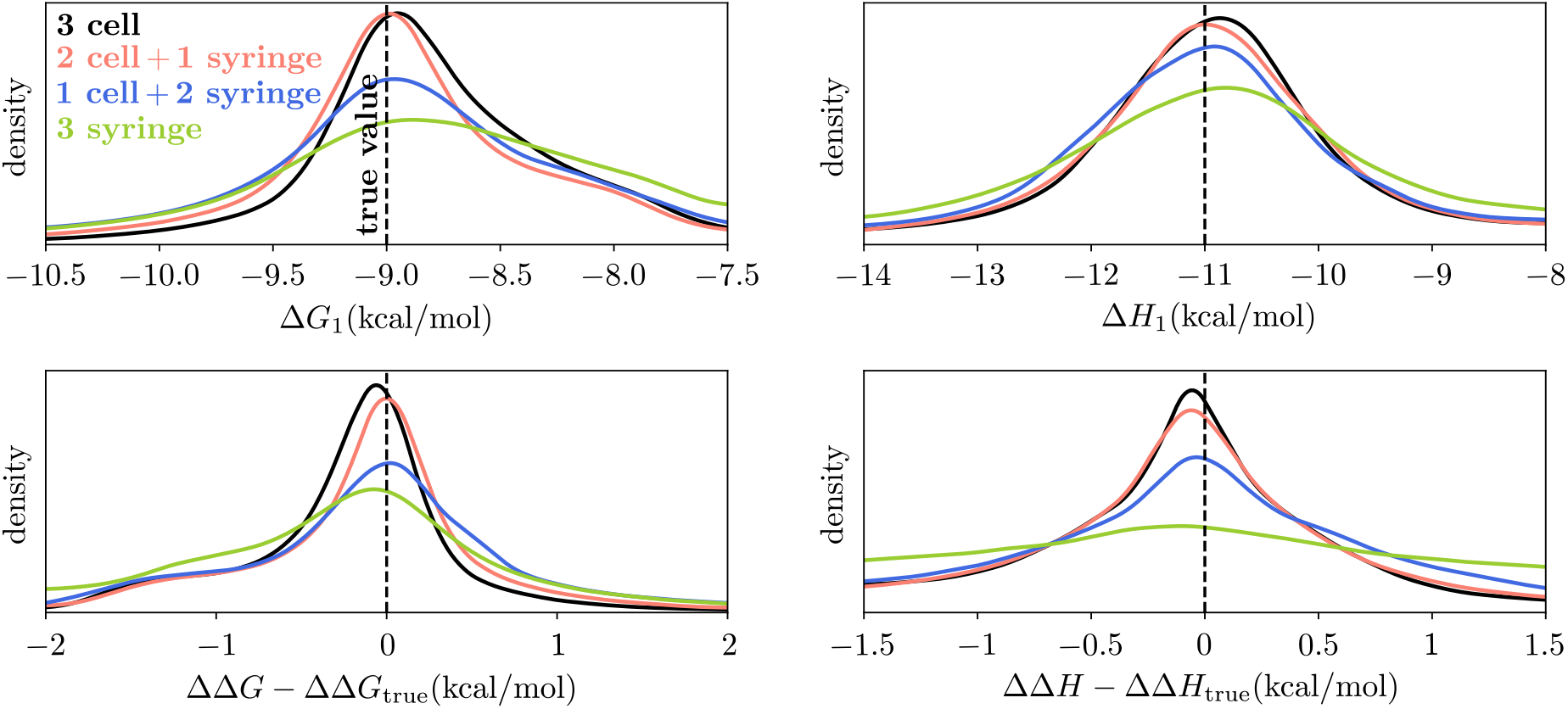
Accumulated posterior marginals for a 2:1 binding model across nine sets of thermodynamic parameters featuring the dimer in the cell 3, 2, 1, or 0 times and in the syringe otherwise. Each accumulated curve is based on the evaluation of 243 independent sets of three synthetic isotherms. The most narrow fits are achieved for the sets featuring the dimer in the cell three times.

**Figure S2:**
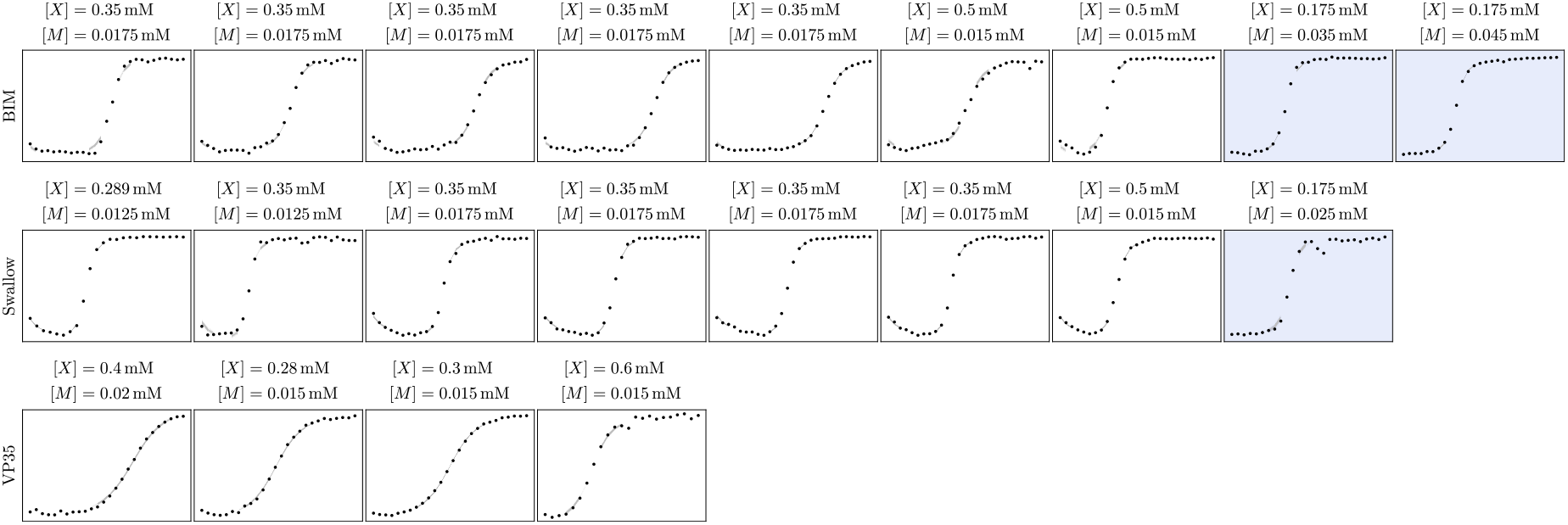
Isotherms of cooperative dimeric binding between the hub protein LC8 and its binding partners Swallow at 10 °C, BIM at 10 °C, and VP35 at 25 °C. Each panel shows the measured heats (points) as well as 1000 isotherms (lines) sampled from the posterior distributions shown in Fig. 8, illustrating the range of fits consistent with the data. The shaded isotherms correspond to experiments in which LC8 was placed in the syringe; all other isotherms were recorded with LC8 in the cell.

## Notes

https://github.com/ZuckermanLab/Bayesian_hierarchical_ITC

### Competing Interest Statement

The authors have declared no competing interest.

